# Multimodal Label-free Monitoring of Adipogenic Stem Cell Differentiation using Endogenous Optical Biomarkers

**DOI:** 10.1101/2020.08.12.246322

**Authors:** Nishir Mehta, Shahensha Shaik, Alisha Prasad, Ardalan Chaichi, Sushant P. Sahu, Syed Mohammad Abid Hasan, Fabrizio Donnarumma, Kermit K. Murray, Ram Devireddy, Manas Ranjan Gartia

## Abstract

Stem cell-based therapies carry significant promise for treating human diseases. However, clinical translation of stem cell transplants for effective therapy requires precise non-destructive evaluation of the purity of stem cells with high sensitivity (< 0.001% of the number of cells). Here, we report a novel methodology using hyperspectral imaging (HSI) combined with spectral angle mapping (SAM)-based machine learning analysis to distinguish differentiating human adipose derived stem cells (hASCs) from control stem cells. The spectral signature of adipogenesis generated by the HSI method enabled identification of differentiated cells at single cell resolution. The label-free HSI method was compared with the standard methods such as Oil Red O staining, fluorescence microscopy, and qPCR that are routinely used to evaluate adipogenic differentiation of hASCs. Further, we performed Raman microscopy and multiphoton-based metabolic imaging to provide complimentary information for the functional imaging of the hASCs. Finally, the HSI method was validated using matrix-assisted laser desorption/ionization-mass spectrometry (MALDI-MS) imaging of the stem cells. The study presented here demonstrates that multimodal imaging methods enable label-free identification of stem cell differentiation with high spatial and chemical resolution. This could provide a powerful tool to assess the safety and efficacy of stem cell-based regenerative therapies.

## INTRODUCTION

Mesenchymal stem cells (MSCs) can be sourced from bone marrow [1], umbilical cord [2], and adipose tissue [3]. Human adipose derived stem cells (hASCs) have been well established as a potential tool for tissue engineering and regenerative medicine applications due to their easy access and lack of ethical issues associated with embryonic stem cells. The hASCs properties of self-renewal, multipotency with respect to differentiation along various cell lineages, and the ability to secrete cytokines and chemokines have been extensively reported during the past decade [4–7]. Adipogenic stem-cell culture requires differentiation times ranging from 9 to 21 days and is expensive. Analysis methods using destructive and invasive techniques such as chemical staining assay, real-time quantitative polymerase chain reaction (qPCR), western blotting assay, and immunostaining to assess the stem cells prevent the re-use of priceless differentiated cells after analysis as these methods require cell lysis, chemical staining, and cell fixation [8]. Label-free non-invasive techniques will facilitate researchers to investigate the cells before clinical transplantation as a quality check and allowing them to perform temporal analysis, thus maximizing the chance of the cell transplant. Hence, label-free approaches capable of probing the stem cells in their native state are required.

Another concern for the clinical translation of hASCs is the formation of teratomas – where a small number of undifferentiated stem cells after transplantation *in vivo* may lead to tumor formation [9, 10]. It is estimated that ~10^4^ number of undifferentiated cells are sufficient to produce teratomas when implanted *in vivo* [10]. Also, for effective treatment, an implantation of about ~ 10^9^ cells is required [11]. That means, an effective method should be capable of discerning differences in cell at a level of 1 in 10^5^ cells (0.001%). Flow cytometry is capable of identifying undifferentiated cells in a cell population at a rate of >1 in 10^3^ (> 0.1%) [12]. Therefore, the sensitivity of flow cytometry is not sufficient to evaluate stem cell purity for transplants. The gold standard method for checking the purity of cells is to transplant them into SCID mice to check for tumor formation [13–15]. But this method takes about 3 months, is expensive, cells might become non-retrievable, and is not quantitative. Therefore, a robust, low-cost, sensitive, and fast method of detecting differentiated stem cells will provide enormous advantages in stem cell therapy.

Previously, several label-free approaches such as electrochemical impedance spectroscopy (EIS)[16–18], Fourier transform infrared spectroscopy (FTIR)[19], Raman spectroscopy[20], and coherent anti-stokes Raman spectroscopy (CARS)[21] were utilized to investigate the progression of stem cell differentiation. Although promising, EIS measurements cannot clearly identify the rate of differentiation in stem cells as there are no significant changes in the impedance signals between cell membrane and electrodes during the progression of differentiation [22]. Due to strong absorption of IR light in water, FTIR spectra of cells maintained under physiological conditions can be distorted and are challenging to carry out. Further, the spatial resolution of FTIR (> 5 μm) is typically smaller than that of Raman imaging (~ 250 nm). Thus, FTIR cell imaging lacks singlecell resolution, and requires complex sample preparation that involves sample drying or fixing [23]. In addition to being expensive, CARS microscopy is limited by capturing single vibrational mode at a time. Further, due to the mixing of nonresonant background (contribution from χ^(3)^) with the signal of resonant vibrational peaks, CARS has limited sensitivity in detecting weak Raman bands. Additionally, the CARS signal has a quadratic relationship with the concentration of the molecules. Hence, it is difficult to detect molecules with low concentrations using CARS [23]. Raman imaging can provide the spectroscopic signatures of the component of stem cells (nucleus, protein, lipids, cytoplasm etc.) with high resolution and specificity[24]. Raman spectroscopy has been used to track adipogenesis [8, 21, 25–27], osteogenesis [8, 28–31], and other stem cell linages [24, 32–37]. However, precise identification of changes in the chemical cues during stem cell differentiation from the Raman spectra obtained from the cellular components (e.g. cytoplasm, nucleus, proteins) are challenging. This is because, significant changes in the cellular components are not expected during stem cell differentiation [8] while significant changes in structure are expected. During Raman analysis of stem cells, auto fluorescence of the cellular components and supporting cell substrates in the fingerprint region obscure the Raman peaks. In addition, photodamage due to laser is a limitation for long-term Raman imaging of the cells [38].

In this paper, we demonstrate a hyper spectral imaging (HSI) method to non-invasively probe the adipogenic differentiation in hASCs (**Figure 1**). The HSI spectral signature is based on the fact that the scattering and absorption of light within stem cells is a function of morphological and molecular changes occurring during the differentiation process[39]. We collected more than fifteen millions scattering spectral signatures using HSI and analyzed using the spectral angle mapping (SAM)-machine learning method to build a library that provided “signature” of adipogenesis. The technique was successfully demonstrated to distinguish differentiated stem cells from control stem cells at an early stage. The HSI results were compared with traditional cell characterization methods (Oil Red O, qPCR), multiphoton imaging, and Raman spectroscopy. Further, the HSI results were validated by performing MALDI-MS imaging of the differentiated and control stem cells. The multimodal imaging approach using HSI and other spectroscopic methods implemented here provides a new tool for label free assessment of stem cells.

**Figure 1.**
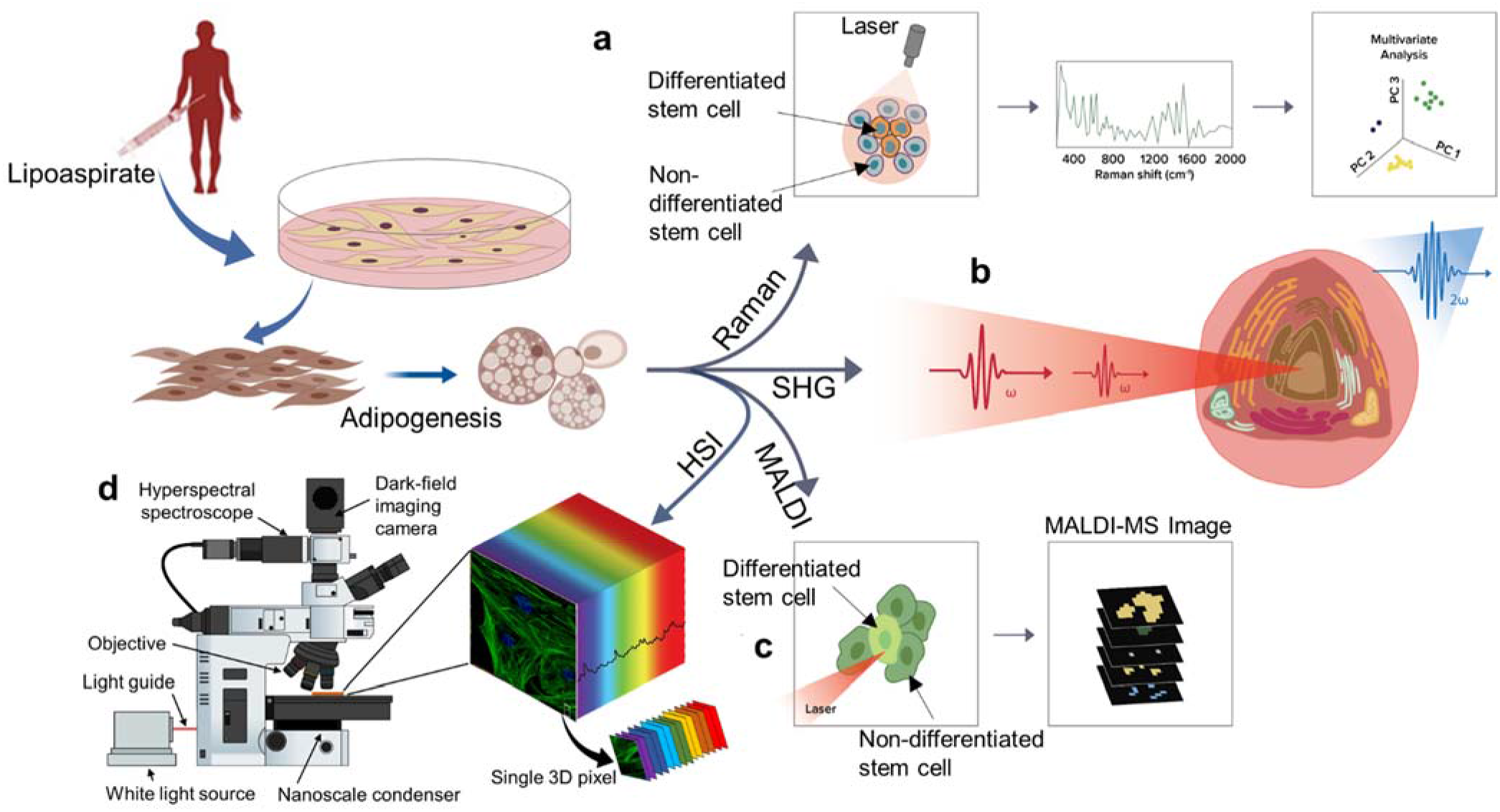
Schematic overview of quantitative label-free imaging implemented at single-cell level. **a.** Label free monitoring of human adipose derived stem cell (hASCs) differentiation to adipogenic stem cells using Raman spectroscopy; **b.** Metabolic imaging using second harmonic generation (SHG) of stem cells to detect differentiated stem cells from control cells; **c.** MALDI-mass spectrometry imaging of lipid in differentiated stem cells; **d.** Schematic arrangement showing the hyperspectral imaging (HSI) system. The HSI system captures the spatial and spectral signature at each pixel of the image to form the data cube.

## MATERIALS AND METHODS

### hASCs Isolation

All the cell-isolation and culture materials were obtained from Sigma-Aldrich (St. Louis, MO) and Fisher Scientific (Hampton, NH). Cell culture protocols used here were approved by the Pennington Biomedical Research Center Institutional Research Board or the Western Institutional Review Board. The cells were isolated from lipoaspirated tissues from unidentified patients with informed consent. Within 24 hours of surgery, the adipo-tissues were thoroughly washed with phosphate-buffered saline (PBS) solution to remove white blood cells (WBCs) and erythrocytes. The PBS solution were warmed to 37 °C and the washing steps were repeated four times. Detailed cell isolation and culture processing procedures are available elsewhere [7, 40]. After isolation, the cells were centrifuged at 300g for 5 min. at 25 °C to separate remaining collagenase material. Finally, the cell pellets were cultured in stromal medium at 37° C, 5% CO_2_, and 95% relative humidity.

### hASCs Differentiation

Adipogenesis was attained by incubating the cells with adipogenic inductive media (ADIPOQUAL™, LaCell LLC, LA, USA). The adipogenic induction media was replaced every 3 days for 10 days. The adipogenic induction media contains IBMX, rosiglitazone, pantothenate, biotin, bovine insulin, penicillin, and dexamethasone. The stromal media used for control samples contains DMEM with 1% antibiotic, and 10% FBS.

### RNA isolation qPCR

RNAs from the stem cells were obtained employing PureLink RNA Mini Kit (Thermo Fisher Scientific, MA, USA), and quantified using a Nanodrop spectrophotometer. cDNA was synthesized using High-Capacity cDNA Reverse Transcription Kit (Thermo Fisher Scientific, MA, USA). Real-time quantitative polymerase chain reaction (qPCR) was performed on an ABI-7900 qPCR machine using the SYBR Green Real-Time PCR Master Mixes (Thermo Fisher Scientific, MA, USA) kit. The primer sequence for the adipogenic genes and cyclophilin B were taken from literature [41]. The fold change in the gene expression was obtained by normalizing it with the housekeeping gene cyclophilin B and quantified using 2^−ΔΔC^_T_ method [42].

### Oil Red O Staining

hASCs cultured with and without adipogenic media at different stages (3, 6, and 9 days) were washed in PBS and were fixed in 4% paraformaldehyde for 30 min at room temperature. After fixation, the PBS rinse step was repeated one more time followed by treatment with 60% isopropanol for 5 min. which is then aspirated, followed by incubating 15 min with the Oil Red O solution. A working solution of Oil Red O prepared with 3 parts of Oil Red O stock solution (0.5% in isopropanol) and 2 parts of distilled water made fresh was used for the staining. The Oil Red O stained cells were washed 2-3 times with PBS and the optical images were captured at 10x magnification and analyzed using ImageJ software. After thresholding for the desired color and hue in ImageJ, images were analyzed based on area and intensity after selecting region on interest (Figure S1-S3).

### Fluorescence Imaging

The cultured control and differentiated hASCs at different stages (0, 3, and 6 days) were fixed in 4% paraformaldehyde, permeabilized with 0.2% TritonX-100 in PBS for 5 min, blocked with 0.2% fish skin gelatin (FSG), and stained with phalloidin (Ex: 488/Em: 510 nm), and counterstained with Hoechst (Ex: 460/Em: 490 nm) for fluorescence imaging. hASCsfor day 9 were fixed, permeabilized, blocked, and stained with Nile red (Ex: 530/Em: 585 nm), and counterstained with Hoechst (Ex: 460/Em: 490 nm). Phalloidin stains the cytoskeletal/actin while Nile Red acts as marker for the mature adipocytes[43]. The concentration of the dye solution was 1 μg/mL and the incubation time was 15 minutes at room temperature. The imaging were performed with a confocal fluorescence microscope (Leica TCS SP8, 63x /1.20 NA water immersion), and analyzed using ImageJ.

### Dark-field Hyperspectral Imaging (HSI)

For the HSI experiments, a CytoViva Hyperspectral Microscope with spectral imaging modality in the visible near-infrared range (λ = 400-1000 nm) was used. The high intensity halogen light source is focused at oblique angles through the liquid light guide for efficient illumination for high contrast imaging in dark field mode. A spectral resolution of 2 nm and spatial pixel width of 25 nm was achieved using the dark-field system. Oil-objectives with 100x magnification were used in our experiment in order to obtain high resolution images. Polarizing filter was placed between condenser and glass substrate in order to minimize random glare. Stem cells were directly grown on an ultra-cleaned glass substrate (Schott) and a coverslip was placed on the cells (fixed in place using nail polish) before imaging. Exposure time for each pixel was set to 0.30 sec. The intensity of the hyperspectral spectrograph was adjusted between 1000 and 10000 counts in order to minimize noise, and avoid saturation of the images. Images were processed using ENVI 4.8 software (CytoViva) in dark-field mode [44]. Spectral information of control and differentiating hASCs obtained from experiments were normalized to 1 for analysis. In the hyperspectral images, each pixel contains the spectrum as a vector. The wavelength range or the spectral bands represent the dimension of the vector. The SAM algorithm compares the vector of each pixel for the control and differentiated stem cells with that of vector of endmember spectrum for classification.

### HSI data processing

Data preprocessing mainly consisted of registration of image, back ground subtraction, and dataset normalization. Normalization of data (with respect to highest intensity in the VNIR spectrum) converted the radiance of HSI observations into reflectance or absorbance [45]. In order to compare each pixel in the selected field of view (FOV), the spectral angle mapper (SAM) algorithm was employed. The SAM algorithm is not sensitive to the intensity of illumination from halogen bulb. In order to normalize the recorded vectors, dark values of the collected data were removed and radiance values were converted to absorbance before using angle mapping as represented in the following equation [45]:

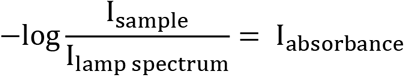

In each pixel of endmember and sampler data cubes, value of n^th^ dimension is the distance between reference origin and the intensity value within that n^th^ band. The best spectral match occurs when the angles between the unit vector of sample image and endmember is the ‘smallest’. In this study, threshold limits of *θ* were restricted to ± 0.10. Therefore, the unknown spectrum is classified as a ‘match’ to the known spectrum if the angles were smaller than the threshold value. The angle within the spectral match was obtained using the following relationship [46]:

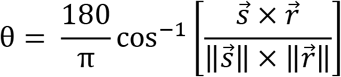

where 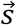 and 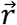 are sample and reference respectively. All the images were calibrated for illumination and reflectance such that the SAM classification was insensitive to albedo or lamp effects. The percentage pixel match data was further weighted based on the area with biomatrix in the FOV.

### Mass Spectrometry Imaging of hASCs using Matrix-assisted laser ionization/deposition (MALDI)

hASCs were grown on indium tin oxide (ITO) coated slides (UniversityWafer Inc, MA, USA). All chemicals for matrix preparation such as methanol, LC-MS grade water, and DHB (2, 5-dihydroxycinnamic acid) were procured from Sigma-Aldrich (St. Louis, MO) while TFA (trifluoroacetic acid) from Fisher Scientific (Atlanta, GA, USA). The matrix solution was prepared with 1:1 ratio of LC-MS grade water and methanol, containing0.1% (v/v) TFA and up to 10 mg/mL of DHB. In order to thoroughly mix solute into solvent, vortexing was employed for 5 min. The slides were vacuum dried for 10 min before matrix deposition. An in-house built pneumatic sprayer with compressed gas (N2) at a pressure of 10 psi was used to spray the matrix at the rate of 50 μL/min on the cells. Approximately 0.5 mL of DHB matrix was used to deposit a uniform layer (spray plus dry cycles coating) over the cells fixated on the ITO slides. Immediately after DHB deposition the slides were heated under a lamp for 10 min until the liquid matrix crystallized. MALDI-MS imaging spectra were acquired using a MALDI-TOF/TOF mass spectrometer (UltrafleXtreme, Bruker Daltonics, Billerica, MA, USA) with 5 ppm mass accuracy. The system was calibrated externally using well-known 5-peptide mix drop cast on the sample slide with 1:1 ratio by volume to matrix. Optical bright field and dark-field images of the samples were taken before matrix deposition for proper calibration via FlexImaging software (Bruker). Each spectrum was produced from 25 laser shots per spot within selected regions of interest (ROI) with step size of 10 μm. Laser shots were fired in spot-array over the 0.05 mm^2^ area.. Spectra were recorded in the positive ion mode in the mass range 400-2,000 *m/z*. MALDI-MS imaging spectra were evaluated using Bruker Flex Analysis 3.0 software and MSiReader (open-source Matlab package) [47, 48] after baseline subtraction using a B-spline fit.

### Multiphoton Microscopy of hASCs

The two-photon fluorescence (TPF) and second harmonic generation (SHG) images of the hASCs were obtained using a Leica SP5 resonant scanning multiphoton confocal microscope with a Spectra Physics Mai-Tai femtosecond tunable pulsed near-IR laser (690 - 1040 nm). A 63x oil objective was used for the image acquisition. The endogenous fluorescence from nicotinamide adenine dinucleotides hydrogen (NADH) and flavin adenine dinucleotides (FAD) [49, 50] were generated by using a Ti:Sa femtosecond laser, with 4000 MHz repetition rate, and 70 fs pulse duration. For NADH, we used an excitation wavelength of 750 nm, and collect emission using filter of 320 – 430 nm. For FAD, we used excitation wavelength of 860 nm, and emission filter of 486 – 506 nm. The redox ratio of FAD/ (NADH+FAD) were calculated at each pixel using ImageJ analysis [51].

### Raman Imaging of hASCs

The Raman spectra and images of the hASCs were obtained using a Raman microspectroscope system (Renishaw inVia Reflex Raman Spectroscope, UK) at a magnification of 50x (long working distance). Laser excitation of 785nm was used with an exposure time of 20s in the wavenumber range of 200–3200 cm^−1^ for acquiring the spectra. For the Raman maps, we used Map Image Acquisition mode of the system. For the Raman images, the following configuration was utilized: grating of 1200 l/mm (633/780), integration time of 2s, wavenumber range of 665-1773 cm^−1^, and with the static mode centered on 1250 cm^−1^. All the Raman spectra were calibrated against the silicon peak at 520 cm^−1^.

### Statistical Analysis

For calculating the *P*-value (statistical significance), two-tailed Student’s t-test or one-way ANOVA were used. The principal component analysis (PCA) was performed using Origin (OriginLab, MA, USA). All the data were expressed in terms of mean and standard deviation.

## RESULTS AND DISCUSSION

Here we utilized HSI to perform label-free chemical imaging to distinguish differentiated hASCs from non-differentiated control cells. Three other label-free modalities (Raman, SHG, MALDI-MS) were used to validate the HSI study and to obtain chemically selective information of the various components of the differentiated stem cells (**Figure 1**). The adipogenic potential of the stem cells were confirmed by staining the differentiated cell population with phalloidin (for cytoskeletal/actin), and Hoechst (nucleus) stains (**Figure 2a – 2c**). To probe the adipogenesis we stained the cells with Nile red and Hoechst (**Figure 2d**) stains. The globular lipid droplet structures within the cells are clearly visible on day 9, representing matured adipocytes (**Figure 2d**). This accumulation of lipid droplets are is a hallmark of adipogenic differentiation process[21, 52]. To further confirm the adipogenic lineage, we investigated the gene expression level of adiponectin[53], leptin[54], and PPARg[55], which were found to be associated with the promotion of adipogenic differentiation. The mRNA fold change of these adipogenic genes were calculated and normalized to cyclophilin B gene by 2^−ΔΔCT^ method [56, 57]. All the gene expression levels increased for hASCs cultured for 9 days in adipogenic media compared to 6 days culture. Especially, the expression level of PPARg, a key regulator for adipocyte differentiation[58], is increased significantly (*P* < 0.05) on day 9 compared to day 6 (**Figure 2e**). The increase in expression level of these genes demonstrate the successful induction of adipogenesis and correlate with the growth of lipid droplets in hASCs with the progression of time.

**Figure 2.**
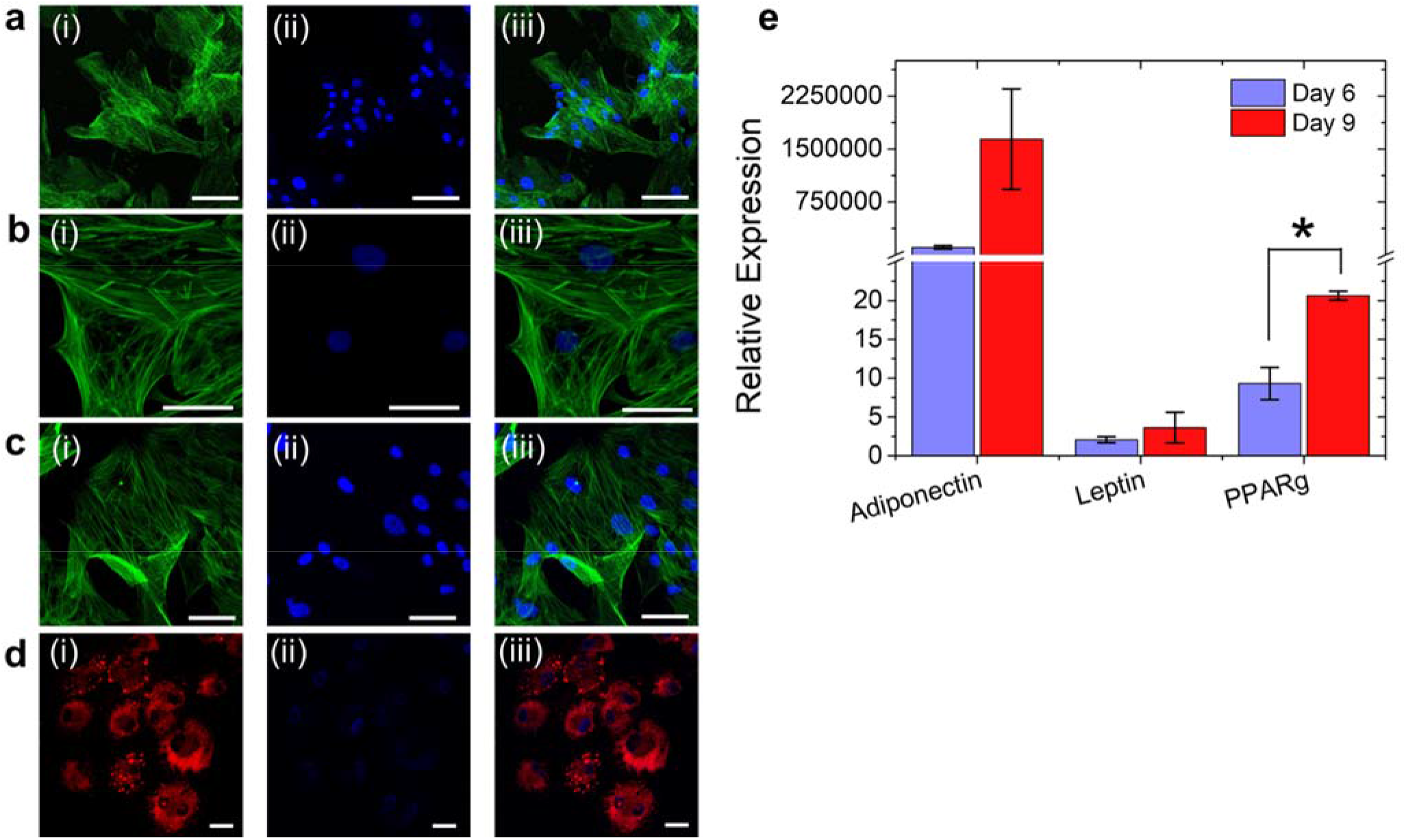
Representative cytochemical image of hASCs and gene expression profile. Confocal fluorescence images of cells cultured in adipogenic media for **a.** 1, **b.** 3, **c.** 6, and **d.** 9 days. The sub-cellular component of each images (scale = 20 μm) are shown for (i) cytoplasm, (ii) nucleus, and (iii) merged. The matured adipocytes (globular lipid droplets) are clearly seen on day 9 with the Nile red stained cells; **e.** Relative gene expression profiles of adiponectin, leptin, and PPARg for stem cells incubated in adipogenic media for 6, and 9 days. Relative expression were normalized to cyclophilin B. The results show the average of three independent donor samples and error bar represent standard deviation. *\*P* < 0.05, calculated using two-tailed Student’s t-test.

The results of conventional histochemical staining of lipids with Oil Red O are shown in **Figure 3**. There is a progressive increase in lipid content with time (day 3 vs. day 9). hASCs cultured in adipogenic media (**Figure 3d, 3e, 3f**) showed significant higher (*P* < 0.001) lipid content compared to the stromal media (**Figure 3a, 3b, 3c**). There was at least a 4-fold increase in the number of droplets on day 9 compared to day 3 for the differentiated cells. These experiments performed over large imaging areas (10x magnification) correlated well with the fluorescence imaging experiments (in **Figure 2**) performed over relatively smaller areas (63x magnification).

**Figure 3.**
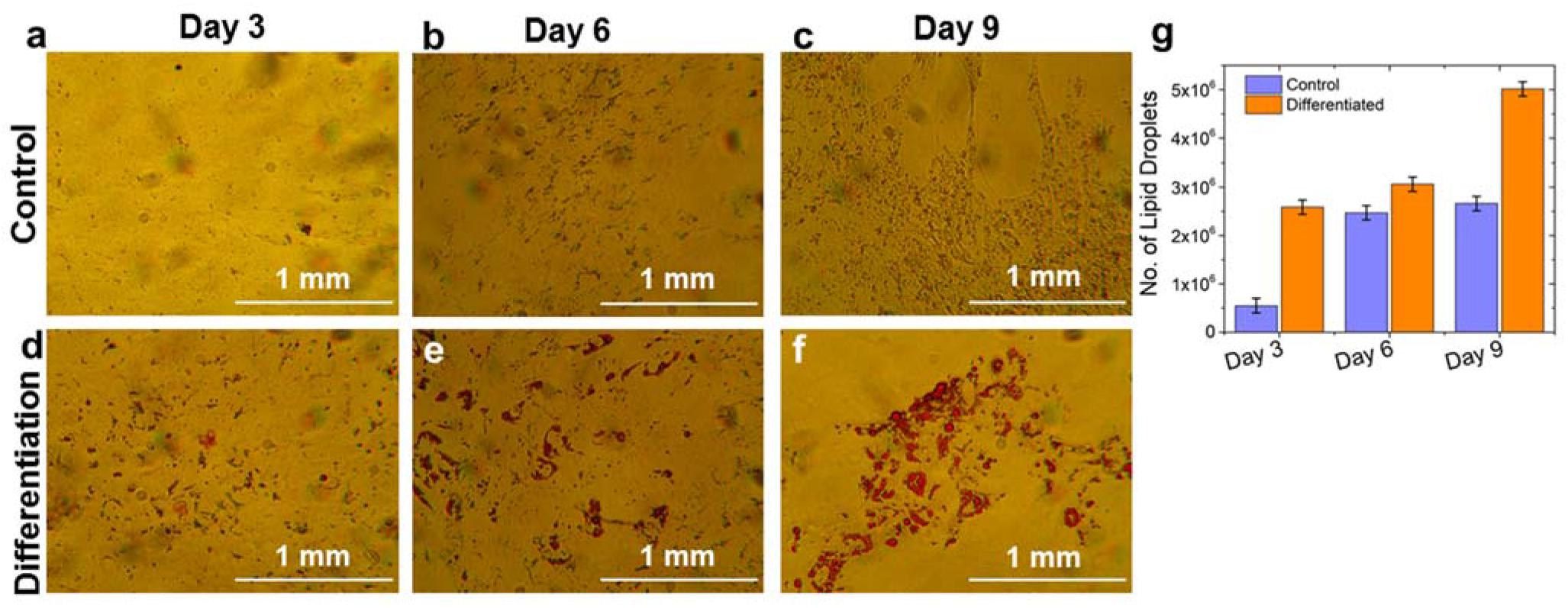
Confirmation of differentiation of hASCs using Oil Red O staining. Comparison of bright field images of Oil Red O staining assay of hASCs for (**a-c**) control (with stromal media), and (d–f) differentiated (with adipogenic media) cells. The cells cultured in stromal and adipogenic media for 3, 6, and 9 days are shown; g. Comparison of number of lipid droplets in differentiated and control cells on day 3, 6, and 9. All the experiments were performed in triplicate. *\*P* < 0.05, \*\**P* < 0.01, \*\*\**P* < 0.001, two-tailed Student’s t-test.

After the conventional validation of success in differentiation of hASCs into adipogenic lineage, we characterized adipogenic differentiation using label-free dark-field microscopy combined with hyperspectral imaging (HSI). Analysis of the acquired dark-field images of biological specimens involves preprocessing of the HSI data cubes, identifying regions of interest and selecting reference spectra of endogenous optical markers to build libraries, and then applying classification algorithms to generate meaningful information. There were a total of 30 field of views (FOVs) from *n* = 3 independent donors captured for both the control and the differentiated samples at days 3, 6 and 9. Each FOV resulted in a generation of 484,416 pixels and in turn each pixel consisting of spatial information along with spectral information in visible near-infrared (VIR) range. That is, the analysis of each image consists of 484,416 spectra and total of 14.5 × 10^6^ spectra for the current study. HSI showed much higher resolution and contrast (approximately 50 nm per pixel) for the lipid droplets without external probes compared to fluorescence and Oil Red O imaging. The stem cells cultured in stromal media (control cells) showed a much smaller number of lipid droplets (**Figure 4a – 4c**) compared to the adipogenic media (**Figure 4d – 4f**). Further, the size and number of droplets in control cells did not change much with time (**Figure 4h**). In contrast, the size of droplets in the differentiated cells progressively grew with time (**Figure 4g, 4h**). Surprisingly, the number of droplets in the differentiated cells decrease with time (**Figure 4h**), possibly due to coalescence of smaller droplets to form larger droplets. The size of the droplets also changed during the differentiation period. For example, on day 3, the majority of the lipid droplets were < 2 μm^2^, whereas on day 9, all of the lipid droplets were > 20 μm^2^ (**Figure 4g**). All of the lipid droplets found in control cells remained < 2 μm^2^ (**Figure 4h**, inset). One of the important advantages of HSI over Oil red O staining and fluorescence imaging is that HSI showed sensitivity towards detecting smaller droplets (early adipogenesis) even on day 3, which are not efficiently imaged using conventional methods (Nile red, Oil red O).

**Figure 4.**
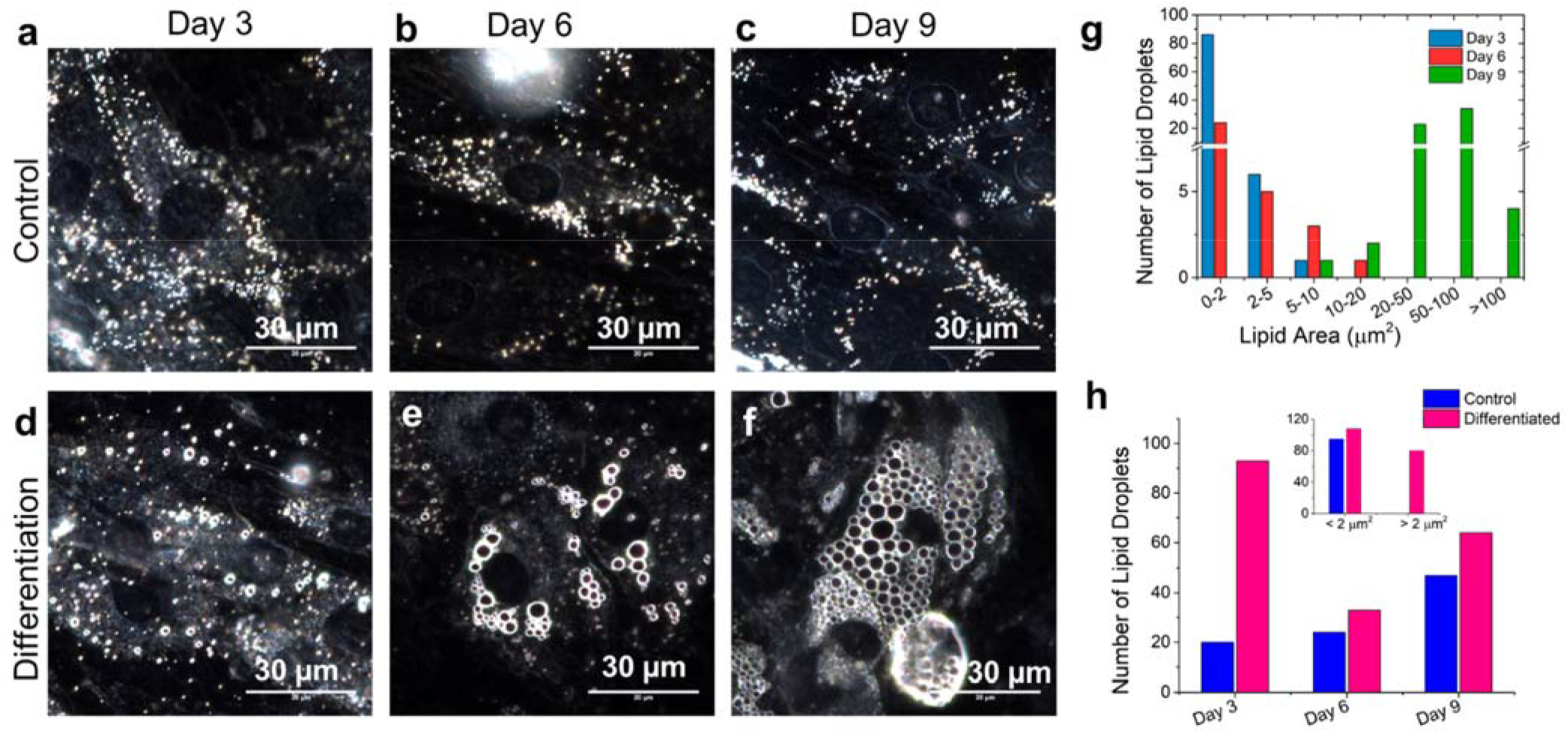
Representative dark-field images showing the morphology of single hASCs cell. Comparison of dark-field images of stem cells in (**a-c**) stromal, (**d-f**) adipogenic media for 3, 6, and 9 days. A progressive increase in the number of larger lipid droplets are visible in the differentiated cells with time compared to control stem cells; **g.** Comparison of size of lipid droplets on different days in differentiated cells. The lipid droplets increased in size withof time; **h.** The distribution of number of lipid droplets for the control and differentiated cells on day 3, 6, and 9. The inset shows that the lipid droplets found in the control cells are primarily of size < 2 μm^2^.

To extract HSI spectral signatures of the differentiated stem cells, we analyzed and found 6 distinct end members (EM) using the SAM method (**Figure 5a**). The characteristic spectral profiles of each endmembers are shown in **Figures 5b - 5g** (each endmember is pseudo colored for visualization). The location of each EMs are shown schematically in **Figure 5h**. EM-1 is originating from the membranes of the lipid droplets (**Figure 5b**) with their spectral peaks at λ = 796 nm. EM-2 (λ = 586 nm) and EM-3 (λ = 721 nm) represent the spectra from the species surrounding the lipid droplets (Figure 5d) and matrix between droplets (**Figure 5c**), respectively. EM-4, with peak at λ = 663 nm, comes from the cytoplasm of the differentiated cell (**Figure 5e**). EM-5 represents the spectra extracted from the fatty esters of oil well since no peak bands were present in the profile (**Figure 5f**). This was an expected outcome due to the refractive index matching of adipogenic oil-wells with microscopy-oil used in the dark-field objective lens. EM-6 (λ = 838 nm) corresponds to smaller lipid droplets (**Figure 5g**). Spectral profiles of EM 1-6 were registered as the reference library in this study and were used in the SAM algorithm. Data presented in **Figure 5i** confirms the spectral distinctiveness of the reference library primary components. One of the most widely used methods to check dataset distinctiveness is principal component analysis (PCA). When PCA is used to check reference library discriminative features, it preserves the variance in high-dimensional space yet reduces redundant information [45] in the bands of HSI images as shown in **Figure 5j**. This further confirms that the HSI spectra result from scattering due to distinctive species and structures within the cells.

**Figure 5.**
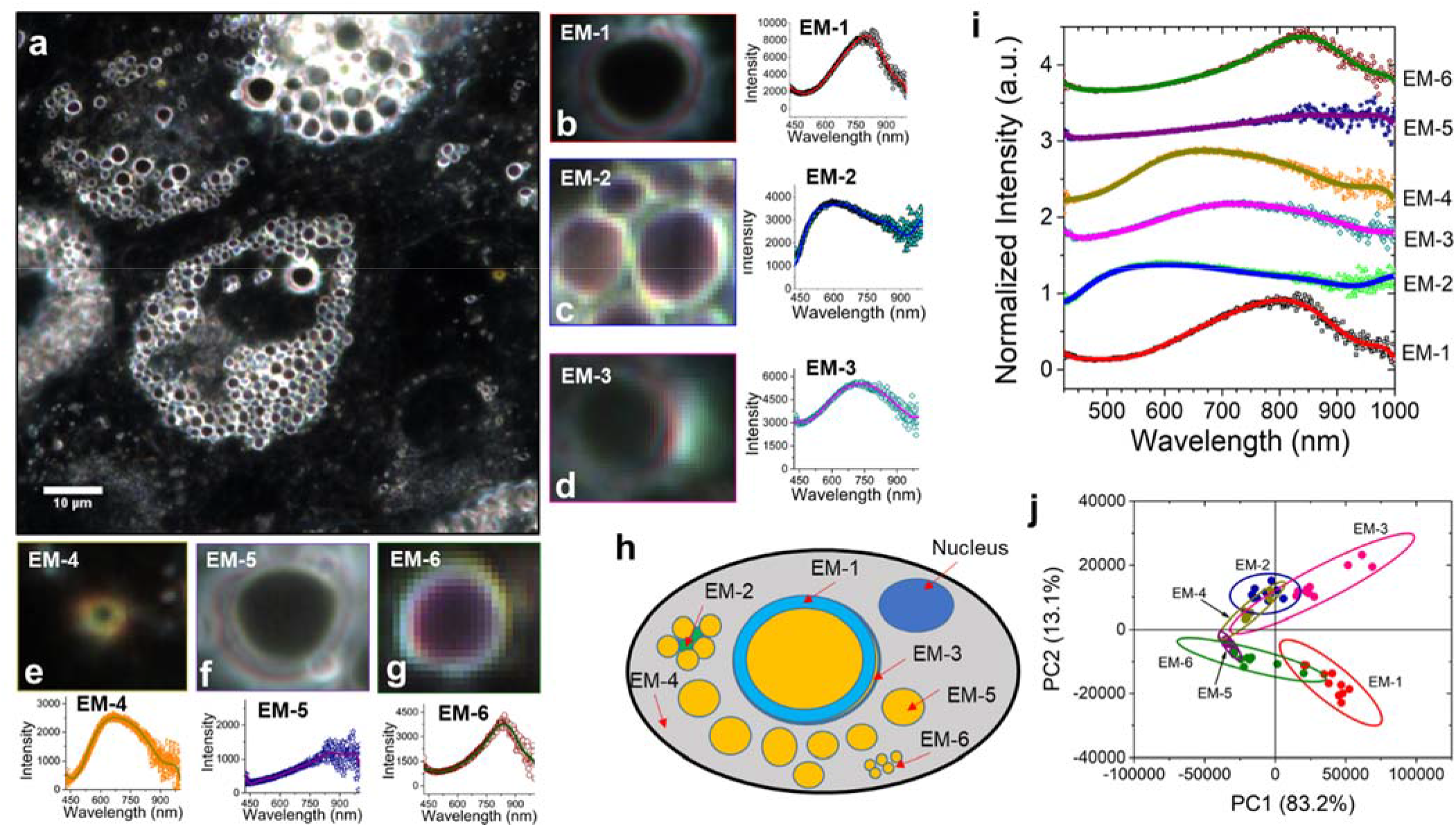
Representative hyperspectral imaging (HSI) of the differentiated hASCs and analysis of their optical biomarkers. **a.** HSI of differentiated stem cells (day 9) with the mapped spectral end members using spectral angle mapper (SAM) analysis. The spectral library was composed of 6 end members (EM-1 to 6). The zoomed in images of each endmembers are shown in **b –g**. The colored area in the images is indicative of the match with the spectral profiles. The spectral signature of the end members in the wavelength range 450 – 950 nm is shown next to the respective end member images; **h.** Schematic figure showing the location inside the cells from which the spectral signature corresponding to each end members is originating; **i.** Comparison of spectral signature of the six end members; **j.** Principal component analysis (PCA) score plot of the six end members for the first two principal components (PC1 vs. PC2). For each endmember, 10 different HSI images were analyzed. Images were acquired using 100x, image size 75 μm x 75 μm, number of samples, n = 3 corresponding to three separate donors.

The spectral library identified earlier was implemented to perform spectral mapping (**Figure 6**) on control and differentiated cells. **Figure 6** provides histograms reporting quantitative percentage classification using the SAM algorithm for each endmember at different time points. The percentage classification describes how many number of pixels in a particular image (out of ~ 500,000 total pixels) belong to a particular endmember. It should be noted that each pixel is composed of spatial as well as spectral information. Among the six spectra, EM-5 represents the main contribution (~ 4%) towards the total scattering of the differentiated cells on day 9 (**Figure 6e**). This is not surprising, since EM-5 is due to the lipid droplets, and by day 9, most of the cell area is occupied by the lipid droplets. It is evident from the histogram that the differentiated stem cells can be clearly distinguished from the control cells onwards of day 6 culture using HSI. The differentiated hASCs on day 9 were found to be significantly different (*P* < 0.001) than the control cells at the same time point while using EM-1 (**Figure 6a**), EM-3 (**Figure 6c**), EM-5 (**Figure 6e**), and EM-6 (**Figure 6f**) classification spectra. On day 6, the analysis showed significant differences (*P* < 0.05) between control and differentiated stem cells for all of the endmembers. The percentage of pixels in the ‘matched’ output on day 3 culture of control and differentiated cells on day 3 was not significant. The percentage classification of endmembers associated with lipid wall periphery (EM-3) and lipid droplets (EM-5) was larger compared to that of other endmembers, which is consistent with lipid droplet formation during the progression of adipogenesis shown earlier (**Figure 3**). Clearly the different rates of cellular metabolism and the differences in the cell morphologies during adipogenic differentiation results in differences in the scattering spectra in comparison to that from control cultures. Thus, HSI provides an alternative quantitative label-free method to monitor hASCs differentiation with minimum sample preparation.

**Figure 6.**
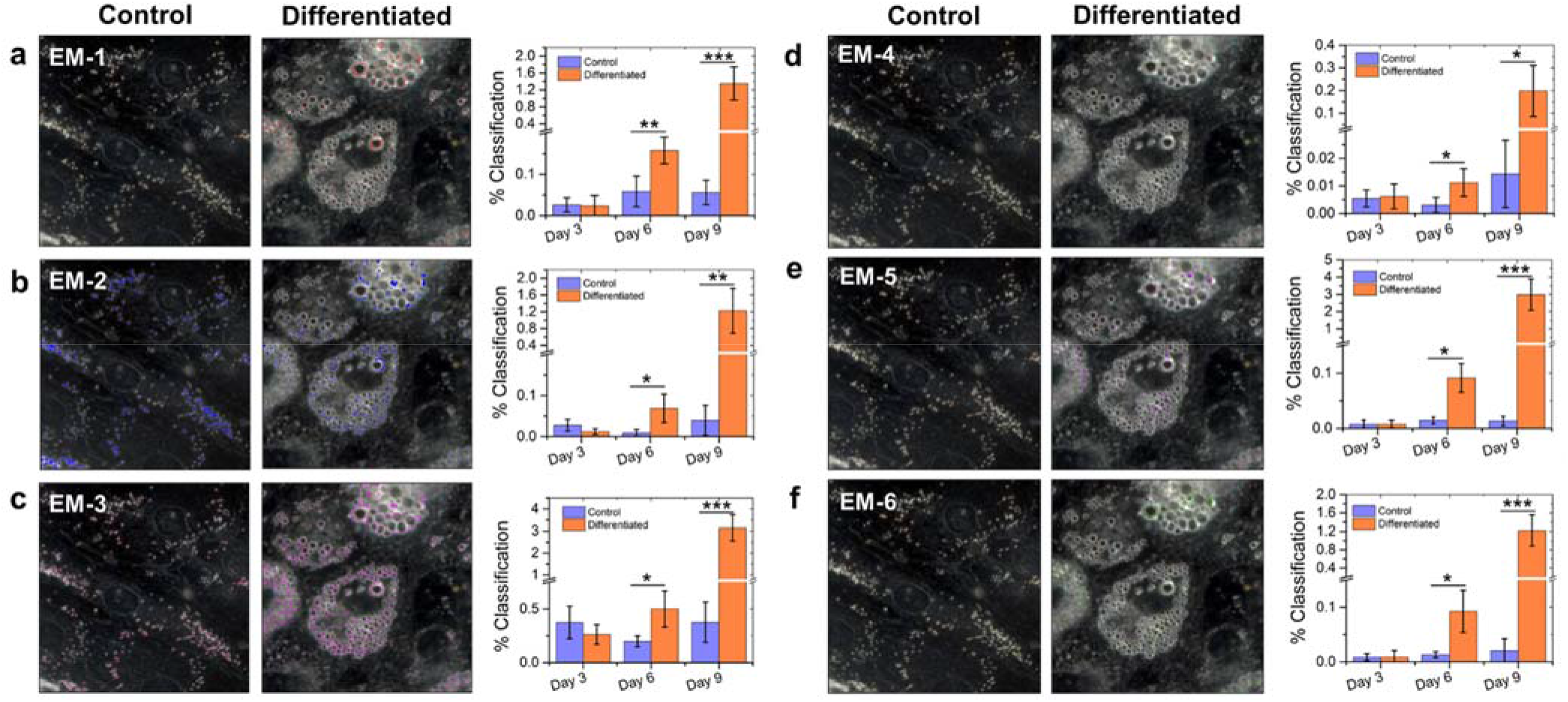
Comparison of Spectral Angle Mapper (SAM) analysis of the control and differentiated hASCs. Representative hyperspectral images on day 9 showing the SAM analysis of control and differentiated cells for **a.** end member 1 (EM-1), **b.** EM-2, **c.** EM-3, **d.** EM-4, **e.** EM-5, and **f.** EM-6. Histogram showing the relative percentage classification (number of matched pixels) for the control and differentiated hASCs on day 3, 6, and 9 are shown for each end members. The number of analyzed images were 15 for each data point and error bar represents the standard deviation. \**P* < 0.05, \*\**P* < 0.01, \*\*\**P* < 0.001, calculated using two-tailed student’s t-test.

To validate the HSI results and identify the molecular origin of each endmembers, we performed MALDI-MS imaging of the control and differentiated hASCs [59, 60]. A comparison of positive ion mode MALDI mass spectra for hASCs cultured for 9 days in adipogenic (differentiated stem cells) and stromal media (control stem cells) is shown in **Figure 7a**. A representative MALDI image of a differentiated cell showing the distribution of FA and DG is shown in **Figures 7b** and **7e**. The corresponding dark-field image of the cells is shown in **Figure 7d**. The higher molecular weight peaks (*m/z* > 1200) seen in the control cell spectrum are absent in differentiated stem cells. Because, we were mainly interested in profiling lipids, we restricted our analysis to lipidomics only. Ganglioside (GD3 d18:1/24:1 *m/z* 1592.881) was not observed in the differentiated stem cells, but is seen in the control cells. Gangliosides are generally responsible for proliferation and transmembrane signaling[61]. They reside in the lipid raft of the membrane[62] (**Figure 7f**) and are found to be lower with adipogenesis[63–65], which is consistent with our observation. In stem cells cultured in stromal media, we found many glycerophospholipids such as phosphatidylcholine (PC 28:2;O2, *m/z* 706.474; PC 34:2, *m/z* 758.573; PC 30:3;O3, *m/z* 786.437), phosphatidylserine (PS 30:1, *m/z* 706.474; PS 35:4, *m/z* 808.459), phosphatidylethanolamine (PE 37:2, *m/z* 758.573; PE O-42:4, *m/z* 832.623), phosphatidic acid (PA O-41:0, m/z 761.644), cardiolipins (CL 70:6, *m/z* 1463.943; CL 74:1, *m/z* 1492.127; CL 72:5, *m/z* 1493.975; CL 78:2, *m/z* 1546.168), and various sphingolipids (ceramide, Cer 44:0;O4, *m/z* 734.66; HexCer 32:3;O2, *m/z* 706.474; gangliosides, Hex(4)-HexNAc(2)-Cer 34:2;O2, *m/z* 1592.891; Hex(2)-NeuAc(2)-Cer 42:2;O2, *m/z* 1592.881). Cardiolipins prevent lipid droplet formation by increasing energy consumption[66]. This could explain why cardiolipins were found in control cells, but not in the differentiated stem cells. Mammalian cell membranes are rich in PC and PE. Alteration in PC and PE can lead to changes in lipid droplet size and their formation[67]. In the MALDI-MS of differentiated cells we observed many hallmarks of lipid droplets such as glycerolipids (TG 52:4, *m/z* 855.747; DG 31:1, *m/z* 553.487; MG O-21:0;O, *m/z* 441.333), sterol lipids (ST 27:1, *m/z* 435.348; ST 28:1, *m/z* 439.333; ST 27:2;O4;S, *m/z* 551.246), and fatty acyls (FA 26:1, *m/z* 465.342). It is well known that di(acyl|alkyl)glycerols (DG) facilitate membrane fusion promoting protein and lipid droplet formation[68, 69]. Tri(acyl|alkyl)glycerols (TG) are also abundantly found inside the lipid droplets (**Figure 7c**). We found PG 40:4 (*m/z* 827.578), a prominent mitochondrial phospholipids, responsible for cell growth and promoter of anaerobic metabolism[70]. Further, the overcrowded adipocytes produce cytokines and adipokines due to stress[71]. This is consistent with the observation of ceramides (Cer 38:0;O3, *m/z* 650.55), which are generally implicated in apoptosis[72]. The full list of lipids found in this study are in the supporting information (Table S1, S2).

**Figure 7.**
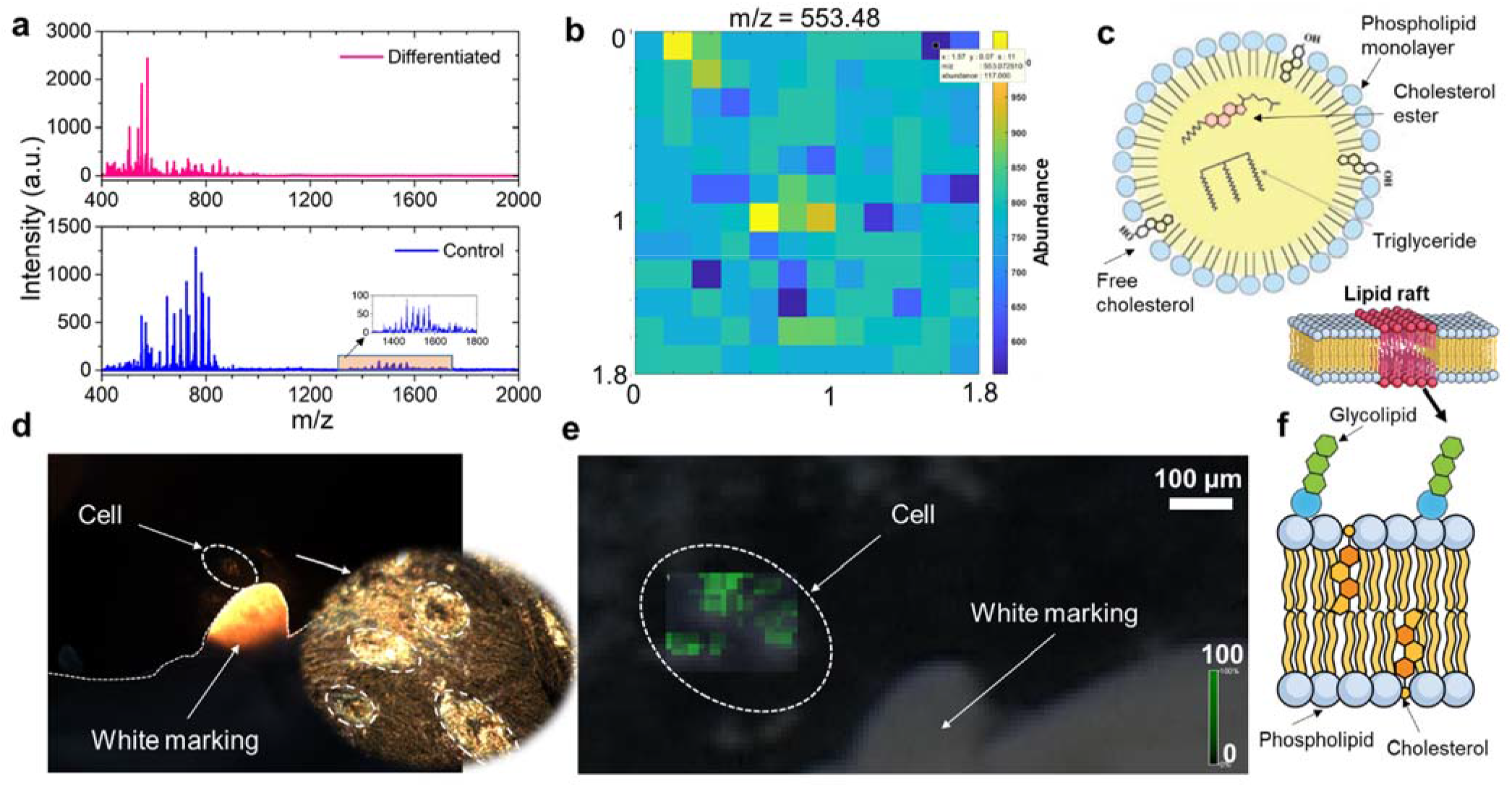
MALDI-MS analysis of hASCs. **a.** Representative MALDI-mass spectra of differentiated and control stem cells on day 9. The inset image shows peaks obtained for the control cells (*m/z* = 1300 - 1800), which are absent in the differentiated cells; **b.** A representative MALDI image of the differentiated cell at *m/z* = 553.48; **c.** schematic showing a lipid droplet and the components of the lipid droplet of the differentiated adipocytes; **d.** dark-field optical image of stem cell on the MALDI substrate. The inset shows the zoomed image of the cells; **e.** MALDI-MS image of the differentiated hADSC shown in d, and overlaid on the respective bright-field optical image; f. schematic showing the cell membrane of a typical differentiated adipogenic stem cell. The lipid raft and the lipids found from the MALDI-MS analysis are shown in the image.

As shown in **Figure 7c**, TGs are abundantly found inside the lipid droplets. TGs in adipose tissue are used for storing energy and synthesis of FAs [73]. Due to the lower oxidation states of carbons of FAs in TGs, it produces more energy than proteins or carbohydrates during oxidative metabolism. Further, due to the highly hydrophobic nature of the TGs, it does not have to carry extra hydrated water as do polysaccharides [74]. To probe the metabolic states of the differentiated hASCs, we performed multimodal (two-photon and SHG) imaging (Figure 8). **Figure 8a – 8c** shows the acquired images of the control cells and **Figure 8d – 8f** showing the images of differentiated stem cells on day 9. We calculated the optical redox ratio (fluorescence intensity ratio of FAD to NADH) to understand the metabolic behavior during adipogenesis [75–79]. We observed a significant decrease (*P* < 0.01) of the redox ratio in the differentiated cells compared to control cells (**Figure 8i, 8j**). Meleshina *et al*. found decrease of the redox ratio (FAD/NADH) during stem cell differentiation[75, 76] which they attributed this to metabolic switching from glycolysis (during undifferentiated state) to oxidative phosphorylation (aerobic metabolism) during adipogenesis[80]. During glycolysis the following reaction occurs:

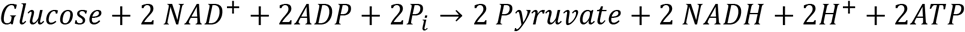

**Figure 8.**
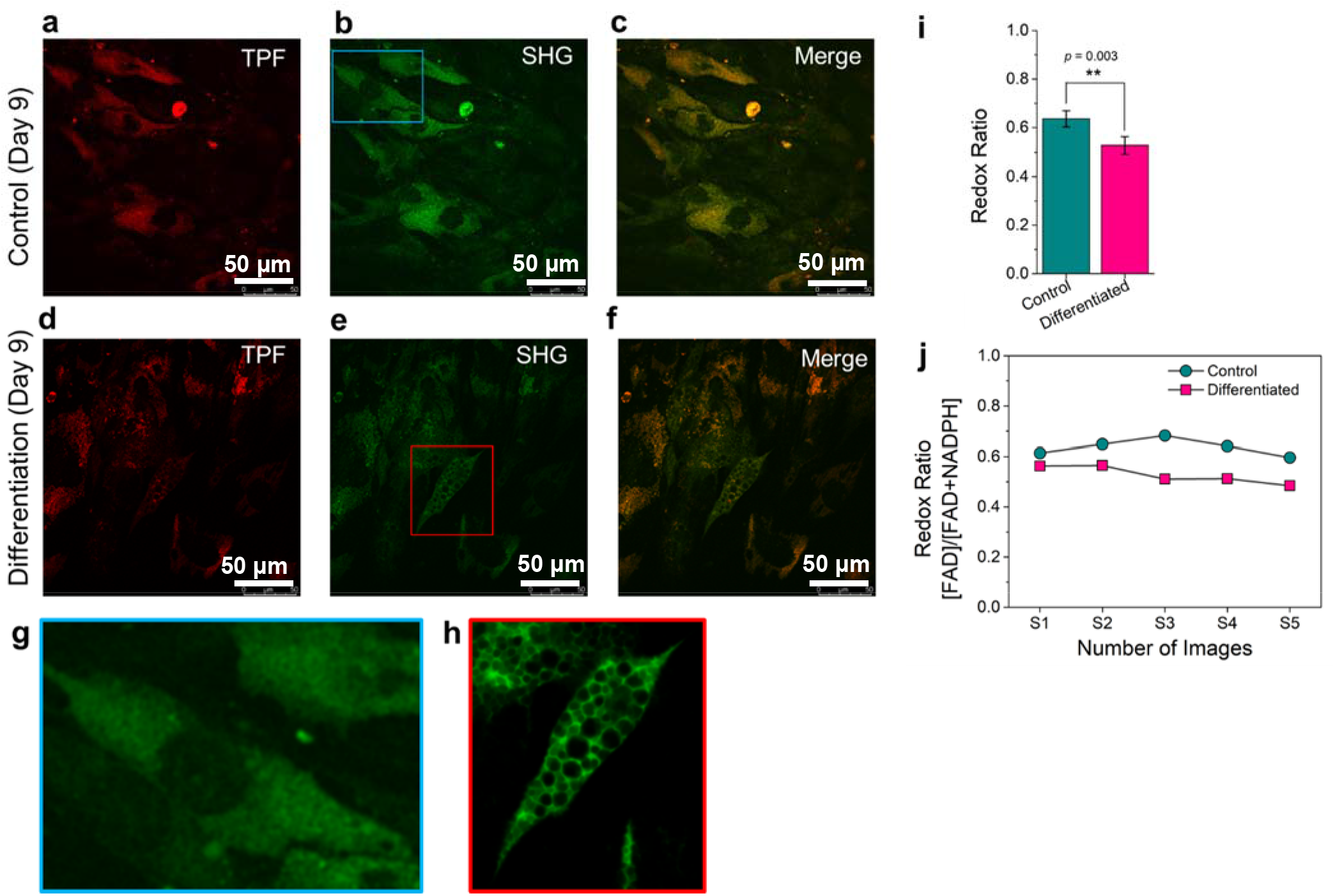
Metabolic imaging of the hASCs using multimodal two-photon fluorescence (TPF) and second harmonic generation (SHG) microscopy. **a.** TPF, **b.** SHG, and **c.** merged image of the hASCs in stromal media on day 9; **d.** TPF, **e.** SHG, and **f.** merged image of the hASCs in adipogenic media on day 9. The zoomed in SHG images are shown for **g.** control cell (blue box in **b**), and **h.** differentiated cells (red box in **e**); **i.** comparison of optical redox ratio [FAD/(NADH+FAD)] between control and differentiated cells; **j.** redox ratio calculated for 5 different images corresponding to control and differentiated hASCs are shown. \*\**P* < 0.01, calculated using two-tailed Student’s t-test.

In this process the FADs are not involved. The number of electrons (and energy produced, 2ATP) in glycolysis is lower compared to oxidative phosphorylation (34 ATP). During oxidative phosphorylation the following reaction occurs:

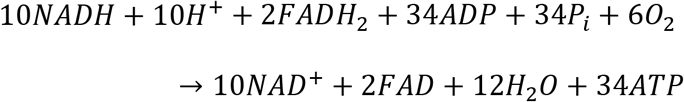

In oxidative phosphorylation, the number of NADH involved are increased and the available FAD are reduced compared to glycolysis leading to decrease of FAD/NADH ratio.

To further investigate the molecular changes during adipogenesis, we obtained Raman spectra of control and differentiated cells (Figure 9). **Figure 9a** compares the average Raman spectrum obtained from control and differentiated hASCs on day 9. Both the control and differentiated cells showed typical vibrational signatures of different cellular components[24, 32, 81, 82] such as proteins (1264, 1304 cm^−1^, N-H, C-H band for amide III; 1654 cm^−1^, amide I/α-helix), amino acids (855 cm^−1^, tyrosine; 874 cm^−1^, tryptophan; 1004 cm^−1^, phenylalanine), nucleic acids (788 cm^−1^, O-P-O stretch, 1098 cm^−1^, PO_2_ stretch), and carbohydrates (1080 cm^−1^, C-O stretch). Here, we focused our analysis to vibrational signature of lipids. We were able to identify three different classes of lipids (TGs, phospholipids, and CEs) using Raman spectroscopy. The characteristic C=O vibration of the ester group (1748 cm^−1^) was seen in the differentiated cells in **Figure 9a** (due to abundant lipid droplets), but not in the control cells. The intensity of vibration due to acyl chains (CH_2_ bending at 1442 cm^−1^ and CH_2_ twisting at 1304 cm^−1^) was greater in the differentiated cells compared to control cells (**Figure 9a**). The peak at 1442 cm^−1^ is also attributed to phospholipids. The vibrational signatures of TGs (855 cm^−1^) due to C-O-O and CH_3_ rocking, and C=O (1748 cm^−1^) were seen in the differentiated cells. The intense peak at 1652 cm^−1^ corresponds to C=C stretching due to unsaturated lipid bonds. The C-C stretch at 1080 cm^−1^ is due to the presence of cholesterol and was found to be higher in differentiated cells due to the efficient formation of lipid droplets. The symmetric stretching mode vibration for =CH_2_ at 2855 cm^−1^ was dramatically larger for the differentiated cells compared to the control due to the lipid droplet formations during adipogenesis (**Figure 9b**). Hence, this vibrational peak at 2855 cm^−1^ can be used to identify lipid droplets[8, 35]. The peak at 2925 cm^−1^ is due to −CH_3_ stretching and corresponds to saturated lipid bond vibration. We calculated the *I*_2855_(*Lipid*)/*I*_2925_(−*CH*_3_) ratio of control cells to be 0.55, and the ratio for the differentiated cells was found to be 1.15. Hence, *I*_2855_/*I*_2925_ can be utilized as a signature of adipogenesis. A significant difference between control and differentiated cells was found in the ratio between cholesterol ester (CE) and free cholesterol (FC), *I*_1748_(*CE*)*I*_721_(*FC*). The *I*_1748_/*I*_721_ ratio for the differentiated cells was 2.86, and for control cells it was 0.33. We did not find significant difference between the unsaturated lipid to saturated lipid ratio for control and differentiated cells. *I*_1652_(*C* = *C*)*I*_1442_(−*CH*_2_) signifies the ratio of unsaturated to saturated lipid content. The *I*_1652_/*I*_442_ for differentiated cells was 0.8 and that of control cells was 0.82. **Figure 9c** shows the representative spectral signature from sub-cellular component of a differentiated (D), and control (C) hADSC on day 9. The corresponding bright field image of the cell is shown in **Figure 9d**. Raman images of the cell at 794 cm^−1^ (nucleic acid, **Figure 9e**), 1002 cm^−1^ (amino acid, **Figure 9f**), 1450 cm^−1^ (lipids, **Figure 9g**), and the merged image (**Figure 9h**) are also shown. Overall, the results demonstrate that the multimodal approach is capable of identifying the stem cell induction to adipogenesis at the single cell level. The developed approach provides a non-destructive, label-free, and cost effective way of analyzing stem cells.

**Figure 9.**
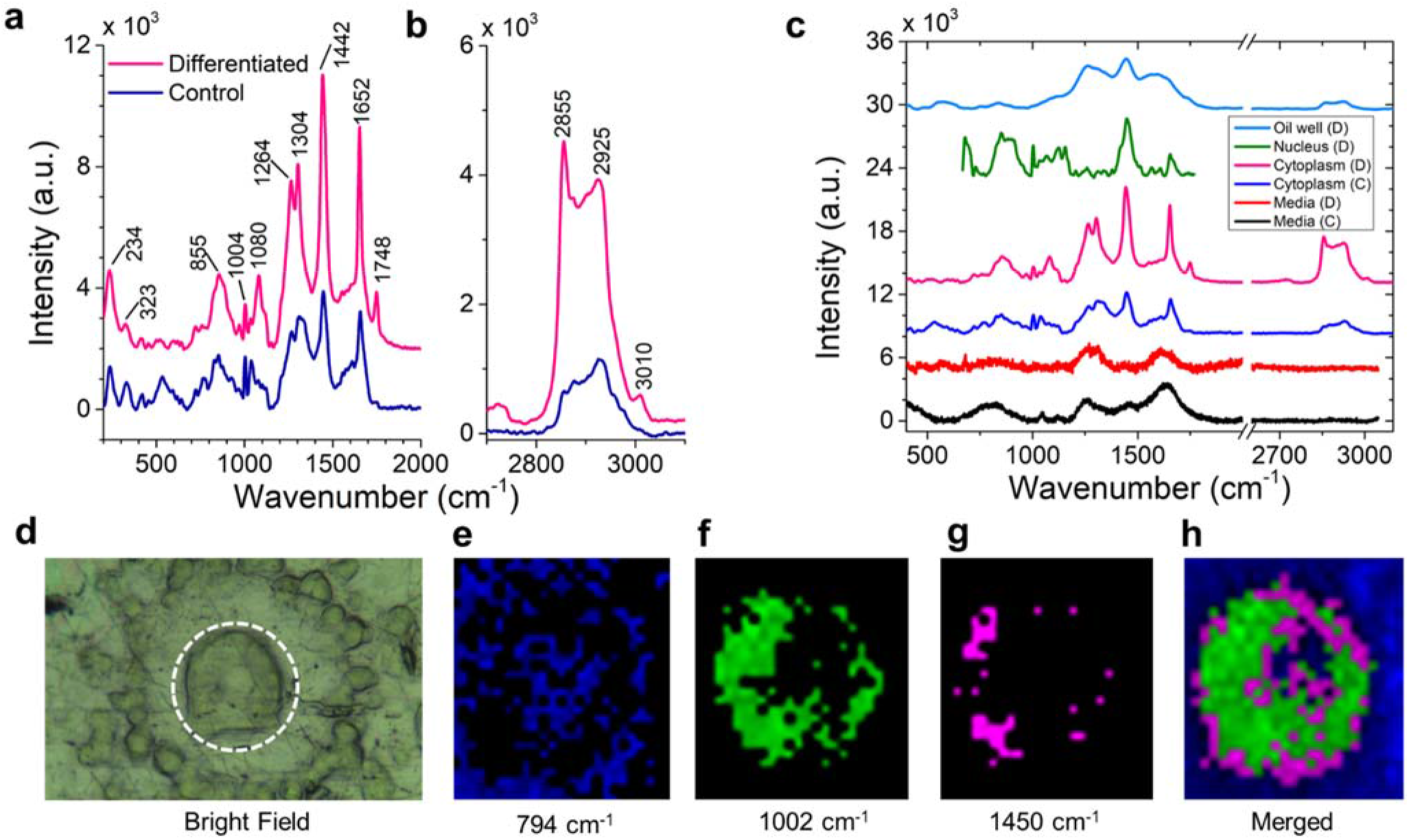
Raman spectroscopy and imaging of hASCs. The average Raman spectra of control and differentiated stem cells from **a.** 400 – 2000 cm^−1^, and **b.** 2700 – 3200 cm^−1^; **c.** the representative spectral signature from sub-cellular component of a differentiated (D), and control (C) hADSC on day 9. The corresponding **d.** bright field image, and Raman image of a single cell at specific vibrational peaks are shown in **e – h**.

## CONCLUSIONS

In this study, we have demonstrated the use of HSI to identify differentiated hASCs from nondifferentiated cells using the endogenous optical and spectral signatures generated from the stem cells. The HSI results were validated using traditional characterization (Oil Red O, qPCR) as well as MALDI-MS, Raman, and multiphoton imaging. HSI method provides better resolution and contrast to observe lipid droplets during adipogenesis compared to conventional Oil Red O staining method. The label-free spectroscopy and chemical imaging methods were shown to be effective in identifying the chemical cues of differentiating stem cells at different days of culture. The results presented here demonstrate that HSI and other label-free method has the potential to provide quantitative molecular imaging information relevant to stem cell biology and therapies that are superior to conventional imaging techniques.

## Acknowledgements

Raman and fluorescence experiments were performed at LSU’s Shared Instrumentation Facility (SIF). SHG experiments were performed at the Genomics or Cell Biology & Bioimaging core facilities that are supported in part by COBRE (NIH8 1P30GM118430-01) and NORC (NIH 1P30-DK072476) center grants from the National Institutes of Health. MRG acknowledges LSU start-up fund and Louisiana Board of Regents Support Fund (RCS Award Contract Number: LEQSF (2017-20)-RD-A-04). AC is supported by LSU Economic Development Assistantships (EDA). Authors thank Dr. David Burk for the help with the fluorescence and SHG experiments.

## Appendix A. Supplementary data

Supplementary data related to this article can be found at https://doi.org/

## Data availability

The authors declare that all data supporting the findings of this study are available within the paper and its supplementary data file. Correspondence and requests for materials should be addressed to M.R.G.

## Notes

### Competing Interest Statement

The authors have declared no competing interest.

