## Supporting information for "Multimodal Label-free Monitoring of Adipogenic Stem Cell Differentiation using Endogenous Optical Biomarkers"

**ImageJ Analysis Steps:**

1. Open Image J (https://imagej.net/Fiji)
2. File→Open→Choose File (Image in .TIF/.JPEG/.JPG)

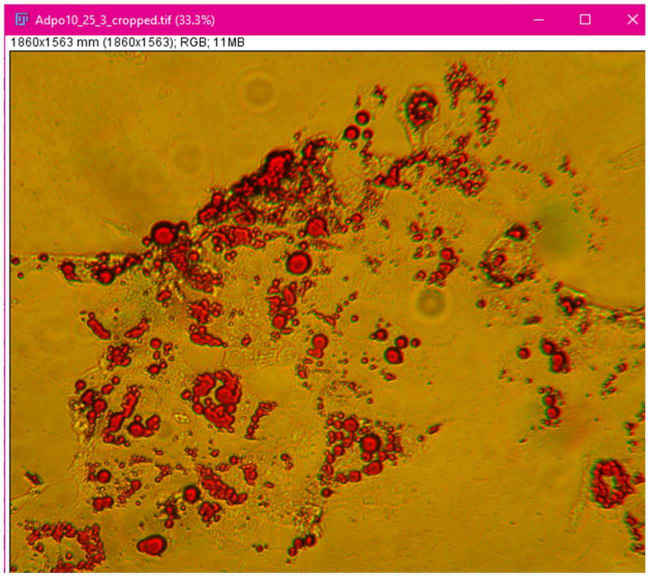


**Figure S1. Original Image**

3. Image→Adjust→Threshold→Color Threshold. In the Threshold Dialog Box, Adjust Hue (Select→ Hue as per desired)

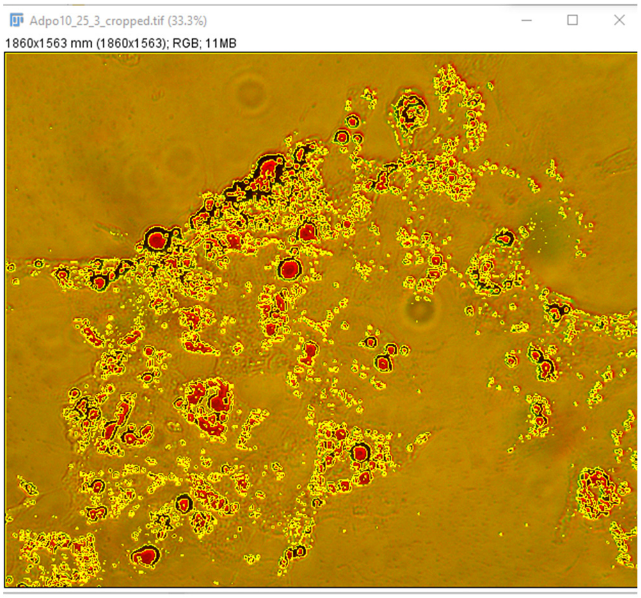


**Figure S2. Thresholding**

4. Analyze →Select ROI (Region of Interest)

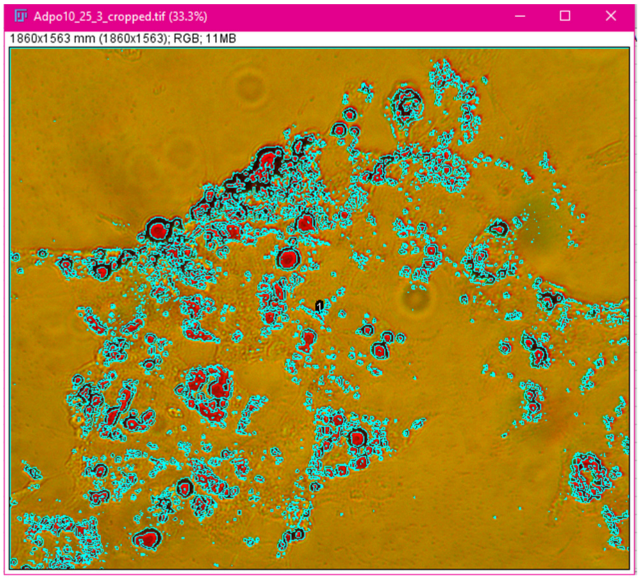


**Figure S3. ROI Highlighted in Blue**

5. Analyze →Set Measurements (Area, Intensity, Minimum, Maximum etc.)→ Measure
6. Save as Measurements Recorded as Excel files.

**Table S1. Matched masses for control stem cells using LIPID MAPS**

**(Positive ion modes, M+H, M+Na, M+K, +/- 0.01 *m/z*)**

Download Matched masses (tab-delimited text)

*This table can be scrolled horizontally if width exceeds container width.*

| Input Mass | Matched Mass | Delta | Name | Formula | Ion | LMSD Examples |
|---|---|---|---|---|---|---|
| 700.567 | 700.5722 | .0052 | HexCer 34:1;O2 | $C_{40}H_{77}NO_8$ | [M+H]+ | Examples |
| 706.474 | 706.4654 | .0086 | PC 28:2;O2 | $C_{36}H_{68}NO_{10}P$ | [M+H]+ | Examples |
| 706.474 | 706.4654 | .0086 | PS 30:1 | $C_{36}H_{68}NO_{10}P$ | [M+H]+ | Examples |
| 706.474 | 706.4655 | .0085 | HexCer 32:3;O2 | $C_{38}H_{69}NO_8K$ | [M+K]+ | Examples |
| 734.66 | 734.6633 | .0033 | Cer 44:0;O4 | $C_{44}H_{89}NO_5Na$ | [M+Na]+ | Examples |
| 758.573 | 758.5694 | .0036 | PC 34:2 | $C_{42}H_{80}NO_8P$ | [M+H]+ | Examples |
| 758.573 | 758.5694 | .0036 | PE 37:2 | $C_{42}H_{80}NO_8P$ | [M+H]+ | Examples |
| 758.573 | 758.5694 | .0036 | PE-NMe 36:2 | $C_{42}H_{80}NO_8P$ | [M+H]+ | Examples |
| 761.644 | 761.6419 | .0021 | PA O-41:0 | $C_{44}H_{89}O_7P$ | [M+H]+ | Examples |
| 761.644 | 761.6531 | .0091 | SM 38:0;O2 | $C_{43}H_{89}N_2O_6P$ | [M+H]+ | Examples |
| 761.644 | 761.6420 | .0020 | DG 43:0 | $C_{46}H_{90}O_5K$ | [M+K]+ | Examples |
| 786.437 | 786.4318 | .0052 | PC 30:3;O3 | $C_{38}H_{70}NO_{11}PK$ | [M+K]+ | Examples |
| 808.459 | 808.4525 | .0065 | PS 35:4 | $C_{41}H_{72}NO_{10}PK$ | [M+K]+ | Examples |
| 832.623 | 832.6191 | .0039 | PE O-42:4 | $C_{47}H_{88}NO_7PNa$ | [M+Na]+ | Examples |
| 837.683 | 837.6820 | .0010 | SM 42:1;O2 | $C_{47}H_{95}N_2O_6PNa$ | [M+Na]+ | Examples |

**Search parameters**

Database: LMSD

Ion adducts: 'M+H', 'M+Na', 'M+K'

Mass Tolerance (m/z): +/- 0.01

Download Matched masses (tab-delimited text)

*This table can be scrolled horizontally if width exceeds container width.*

| Input Mass | Matched Mass | Delta | Name | Formula | Ion | LMSD Examples |
|---|---|---|---|---|---|---|
| 1409.578 | 1409.5832 | .0052 | LPIM5 18:1 | $C_{57}H_{101}O_{37}P$ | [M+H]+ | Examples |
| 1463.943 | 1463.9354 | .0076 | CL 70:6 | $C_{79}H_{142}O_{17}P_2K$ | [M+K]+ | Examples |
| 1492.127 | 1492.1204 | .0066 | CL 74:1 | $C_{83}H_{160}O_{17}P_2$ | [M+H]+ | Examples |
| 1493.975 | 1493.9824 | .0074 | CL 72:5 | $C_{81}H_{148}O_{17}P_2K$ | [M+K]+ | Examples |
| 1546.168 | 1546.1673 | .0007 | CL 78:2 | $C_{87}H_{166}O_{17}P_2$ | [M+H]+ | Examples |
| 1592.891 | 1592.8894 | .0016 | Hex(4)-HexNAc(2)-Cer 34:1;O2 | $C_{74}H_{133}N_3O_{33}$ | [M+H]+ | Examples |
| 1592.891 | 1592.8813 | .0097 | Hex(2)-NeuAc(2)-Cer 42:2;O2 | $C_{76}H_{135}N_3O_{29}K$ | [M+K]+ | Examples |

**Table S2. Matched masses for differentiated stem cells using LIPID MAPS**

| Input Mass | Matched Mass | Delta | Name | Formula | Ion | LMSD Examples |
|---|---|---|---|---|---|---|
| 423.174 | 423.1780 | .0040 | FA 20:4;O5 | $C_{20}H_{32}O_7K$ | [M+K]+ | Examples |
| 433.362 | 433.3676 | .0056 | ST 28:1;O3 | $C_{28}H_{48}O_3$ | [M+H]+ | Examples |
| 433.362 | 433.3652 | .0032 | FA 26:1;O | $C_{26}H_{50}O_3Na$ | [M+Na]+ | Examples |
| 435.348 | 435.3469 | .0011 | ST 27:1;O4 | $C_{27}H_{46}O_4$ | [M+H]+ | Examples |
| 435.348 | 435.3445 | .0035 | FA 25:1;O2 | $C_{25}H_{48}O_4Na$ | [M+Na]+ | Examples |
| 435.348 | 435.3445 | .0035 | FA 26:1;O2 | $C_{25}H_{48}O_4Na$ | [M+Na]+ | Examples |
| 439.333 | 439.3418 | .0088 | ST 26:0;O5 | $C_{26}H_{46}O_5$ | [M+H]+ | Examples |
| 439.333 | 439.3337 | .0007 | ST 28:1;O | $C_{28}H_{48}OK$ | [M+K]+ | Examples |
| 441.333 | 441.3363 | .0033 | ST 29:4;O3 | $C_{29}H_{44}O_3$ | [M+H]+ | Examples |
| 441.333 | 441.3339 | .0009 | ST 27:1;O3 | $C_{27}H_{46}O_3Na$ | [M+Na]+ | Examples |
| 441.333 | 441.3341 | .0011 | MG O-21:0;O | $C_{24}H_{50}O_4K$ | [M+K]+ | Examples |
| 447.359 | 447.3597 | .0007 | ST 30:3;O | $C_{30}H_{48}ONa$ | [M+Na]+ | Examples |
| 447.359 | 447.3599 | .0009 | FA 27:1 | $C_{27}H_{52}O_2K$ | [M+K]+ | Examples |
| 449.377 | 449.3754 | .0016 | ST 30:2;O | $C_{30}H_{50}ONa$ | [M+Na]+ | Examples |
| 449.377 | 449.3755 | .0015 | FA 27:0 | $C_{27}H_{54}O_2K$ | [M+K]+ | Examples |
| 461.239 | 461.2429 | .0039 | LPA O-18:1 | $C_{21}H_{43}O_6PK$ | [M+K]+ | Examples |
| 461.239 | 461.2452 | .0062 | ST 28:6;O3 | $C_{28}H_{38}O_3K$ | [M+K]+ | Examples |
| 461.239 | 461.2300 | .0090 | ST 24:2;O6 | $C_{24}H_{38}O_6K$ | [M+K]+ | Examples |
| 465.342 | 465.3452 | .0032 | LysoSM(d18:1) | $C_{23}H_{49}N_2O_5P$ | [M+H]+ | Examples |
| 465.342 | 465.3339 | .0081 | ST 29:3;O3 | $C_{29}H_{46}O_3Na$ | [M+Na]+ | Examples |
| 465.342 | 465.3493 | .0073 | ST 30:2;O | $C_{30}H_{50}OK$ | [M+K]+ | Examples |
| 465.342 | 465.3341 | .0079 | FA 26:1;O2 | $C_{26}H_{50}O_4K$ | [M+K]+ | Examples |
| 465.342 | 465.3341 | .0079 | FA 27:1;O2 | $C_{26}H_{50}O_4K$ | [M+K]+ | Examples |
| 483.333 | 483.3235 | .0095 | ST 29:2;O3 | $C_{29}H_{48}O_3K$ | [M+K]+ | Examples |
| 501.312 | 501.3211 | .0091 | ST 30:5;O6 | $C_{30}H_{44}O_6$ | [M+H]+ | Examples |
| 545.291 | 545.2874 | .0036 | ST 27:2_O5 | $C_{27}H_{45}O_9P$ | [M+H]+ | Examples |
| 545.291 | 545.2875 | .0035 | ST 29:3;O7 | $C_{29}H_{46}O_7K$ | [M+K]+ | Examples |
| 551.246 | 551.2439 | .0021 | ST 27:2;O4;S | $C_{27}H_{44}O_7SK$ | [M+K]+ | Examples |
| 553.487 | 553.4826 | .0044 | DG 31:1 | $C_{34}H_{64}O_5$ | [M+H]+ | Examples |
| 553.487 | 553.4955 | .0085 | FA 36:3 | $C_{36}H_{66}O_2Na$ | [M+Na]+ | Examples |
| 569.306 | 569.3085 | .0025 | PG 20:1;O | $C_{26}H_{49}O_{11}P$ | [M+H]+ | Examples |
| 599.279 | 599.2746 | .0044 | LPG 22:4 | $C_{28}H_{49}O_9PK$ | [M+K]+ | Examples |
| 650.55 | 650.5484 | .0016 | Cer 38:0;O3 | $C_{38}H_{77}NO_4K$ | [M+K]+ | Examples |
| 677.6 | 677.6078 | .0078 | DG 40:2 | $C_{43}H_{80}O_5$ | [M+H]+ | Examples |
| 734.66 | 734.6633 | .0033 | Cer 44:0;O4 | $C_{44}H_{89}NO_5Na$ | [M+Na]+ | Examples |
| 827.578 | 827.5797 | .0017 | PG 40:4 | $C_{46}H_{83}O_{10}P$ | [M+H]+ | Examples |
| 827.578 | 827.5773 | .0007 | PG 38:1 | $C_{44}H_{85}O_{10}PNa$ | [M+Na]+ | Examples |
| 855.747 | 855.7436 | .0034 | TG 52:4 | $C_{55}H_{98}O_6$ | [M+H]+ | Examples |
| 855.747 | 855.7412 | .0058 | TG 50:1 | $C_{53}H_{100}O_6Na$ | [M+Na]+ | Examples |

[illegible]
